# Machine learning approaches for estimating cross-neutralization potential among FMD serotype O viruses

**DOI:** 10.1101/2024.05.22.594549

**Authors:** Dennis N Makau, Jonathan Arzt, Kimberly VanderWaal

**Affiliations:** Department of Biomedical and Diagnostic Sciences, College of Veterinary Medicine, University of Tennessee, Knoxville, USA; Foreign Animal Disease Research Unit, USDA-ARS, Plum Island Animal Disease Center, Orient Pt., NY 11957, USA; Department of Veterinary Population Medicine, College of Veterinary Medicine, University of Minnesota, USA

**Keywords:** cross-protection, r_1_ values, cross-reactivity, viral neutralization, machine learning, bioinformatics, computational immunology

## Abstract

In this study, we aimed to develop an algorithm that uses sequence data to estimate cross-neutralization between serotype O foot-and-mouth disease viruses (FMDV) based on r1 values, while identifying key genomic sites associated with high or low r1 values. The ability to estimate cross-neutralization potential among co-circulating FMDVs in silico is significant for vaccine developers, animal health agencies making herd immunization decisions, and disease preparedness. Using published data on virus neutralization titer (VNT) assays and associated VP1 sequences from GenBank, we applied machine learning algorithms (BORUTA and random forest) to predict potential cross-reaction between serum/vaccine-virus pairs for 73 distinct serotype O FMDV strains. Model optimization involved tenfold cross-validation and sub-sampling to address data imbalance and improve performance. Model predictors included amino acid distances, site-wise amino acid polymorphisms, and differences in potential N-glycosylation sites.

The dataset comprised 108 observations (serum-virus pairs) from 73 distinct viruses with r1 values. Observations were dichotomized using a 0.3 threshold, yielding putative non-cross-neutralizing (< 0.3 r1 values) and cross-neutralizing groups (≥ 0.3 r1 values). The best model had a training accuracy, sensitivity, and specificity of 0.96 (95% CI: 0.88-0.99), 0.93, and 0.96, respectively, and an accuracy of 0.94 (95% CI: 0.71-1.00), sensitivity of 1.00, and specificity of 0.93, positive, and negative predictive values of 0.60 and 1.00, respectively, on one testing dataset and an accuracy, AUC, sensitivity, specificity, and predictive values all approaching 1.00 on a second testing dataset. Additionally, amino acid positions 48, 100, 135, 150, and 151 in the VP1 region alongside amino acid distance were found to be important predictors of cross-neutralization.

Our study highlights the value of genetic/genomic data for informing immunization strategies in disease management and understanding potential immune-mediated competition amongst related endemic strains of serotype O FMDVs in the field. We also showcase leveraging routinely generated sequence data and applying a parsimonious machine learning model to expedite decision-making in selection of vaccine candidates and application of vaccines for controlling FMD, particularly serotype O. A similar approach can be applied to other serotypes.

## Introduction

Foot and mouth disease (FMD) is a viral disease caused by the FMD virus (FMDV), a member of the *Piconaviridae* family (1) that affects cloven-hoofed ungulates. Though typically not fatal, the impact of FMD on food security and livelihoods in endemic countries (particularly low-and middle-income countries (LMICs)) cannot be overemphasized (2). FMD also continues to be a major stumbling block to livestock production and global trade in many parts of the world, especially with the continued threat of emergence and introduction of new FMDV lineages/strains into FMD-free countries. As such, numerous efforts and pathways to achieve and maintain disease free status in most countries has required, among other things, the ability and capacity to identify viruses and vaccinate animals with appropriate vaccines for countries where elimination is yet to be achieved. Although there are 7 documented antigenically distinct serotypes (O, A, C, SAT1, SAT2, SAT3 and Asia1), serotype A and O have been reported to be the most common causes of FMD globally (3–6).

Continued efforts for disease management have included the identification of suitable vaccine candidates and formulation of vaccines applied to the animal populations at risk. To identify these vaccine candidates, a process known as vaccine matching is done, which entails the use of viral neutralization titer (VNT) assays to identify effective cross-reaction and cross-neutralization of viruses by candidate vaccines. The measure of effectiveness of the cross-neutralization and subsequent cross-protection is expressed as the r_1_ value and is briefly defined as the ratio between the neutralizing serum titer against the heterologous virus (usually a field strain) to the neutralizing serum titer against the homologous virus (usually the vaccine strain). Subsequently, r_1_ values between virus-serum pairs of ≥ 0.3 are considered evidence of cross-neutralization between the two involved viruses, while r_1_ values < 0.3 are considered to be non-neutralizing (7).

Generating r_1_ data for comparison demands significant effort, precision, and laboratory resources. This involves producing sera in live animals and running *in vitro* assays, to evaluate the cross-reactivity of target viruses with various sera samples. Advances in computational biology empower us to harness machine learning and artificial intelligence to build computation tools aimed at estimating the cross-neutralization potential between different viruses. While previous studies have explored predictive models of FMDV antigenicity, they have varied objectives. Some focused on predicting FMDV antigenicity and identifying antigenicity descriptors (8), identifying and predicting specific epitopes and genomic regions of antigenic importance (9, 10), or developing intricate models to forecast antigenic changes among SAT 1 and 2 FMDV serotypes (11–13). There is hardly any literature specific to serotype O despite it being one of two widely occurring serotypes, nor has research on the application of machine learning to estimate r_1_ values been published. Although there have been some divergent views about the adequacy of using r_1_ values in identifying potential cross-protection (14), it is still one of the metrics relied upon when interpreting data generated from *in vitro* VNT assays and selecting candidate vaccine strains. Successful application of machine learning to aid in decision making when selecting vaccine candidates and immunization programs has been demonstrated in diseases such as influenza and dengue virus (15–17). Our intention in this study was to develop a simple, yet robust, predictive tool that can be used to estimate potential cross-neutralization between viruses by leveraging machine learning algorithms and genetic characteristics of FMDV.

Therefore, the objective of this study was to develop an algorithm to estimate potential cross-neutralization between strains of serotype O FMDV using r_1_ values and identify important genomic sites that influence high or low r_1_ values between viruses. The ability to distinguish and estimate the potential for cross-protection between different cocirculating FMD viruses *in silico* will support decision-making by vaccine developers in vaccine candidate matching, animal health agencies/institutions in decision-making for herd immunization practices, and overall disease preparedness and response in cases of emergent strains especially for serotype O about which there is a dearth of information.

## Materials and methods

### Data preparation

We obtained data from published manuscripts on r_1_ values for FMDV serotype O. Viruses utilized in these papers came from 14 countries. These manuscripts were obtained from a search from google and PubMed databases using the search terms ‘FMDV and r_1_ values’ and ‘FMDV and vaccine matching’ yielding 4 manuscripts (18–21). To be included, the study must have followed standard WHOA (22) methodology for generation of r_1_ values for serotype O viruses using virus neutralization tests (VNTs), as well as have associated viral sequence data. The r_1_ values indicate the serological compatibility between the vaccine strain and the field isolate, determined by comparing the reactivity of the heterologous versus homologous virus to the antisera produced against the specific viruses. The ratio of the heterologous to homologous neutralization titers between field isolates and vaccine sera serve as a measure of cross-protection.

Since majority of the published studies had uploaded only VP1 data to GenBank, we downloaded sequence data for the VP1 gene of 73 distinct viruses used in the cross-reactivity and generation of r_1_ data summarized in the manuscripts described above and any whole genomes available were trimmed to the VP1 region using Muscle in Aliview 1.26 (23). Accession numbers to the sequences used in this analysis are available in supplemental material 1. Using Muscle in Aliview 1.26 (23) we aligned and translated sequences into amino acid sequences. Using MEGA X (24), we calculated Poisson corrected amino acid distance between all pairs (24, 25) and used R packages *stringr* (26), *bioseq* (27), *ape* (28) and *tidysq* (29) to code site-wise differences at any polymorphic amino acid sites using R v4.2 software (30). For each polymorphic amino acid site, serum-virus pairs were tabulated as 0 if they shared the same amino acid and 1 if they had a different amino acid at that site. We also scanned the genome for sites of potential N-glycosylation (31, 32) in the VP1 region using a custom-built R script. Pairs with identical sets of inferred glycosylated sites were coded as 0 and non-identical sets as 1. The r_1_ values, as reported in the manuscripts, amino acid distance calculated in MEGA X, and amino acid site-wise differences were concatenated into one data frame, resulting in 108 observations (serum-virus pairs). Although some studies on influenza have explored the pros and cons of weighting of specific amino acid sites based on *a priori* knowledge of their evolutionary history and influence on antigenicity (33, 34), our modeling approach was naïve to *a priori* assumptions about the relative importance of specific site-wise amino acid differences and thus no weighting was used (e.g., I→N ≡ N→I) (15).

Initial analysis involved descriptive statistics of the data and correlation analysis. We performed a phylogenetic analysis of the sequence data by constructing a bootstrapped maximum likelihood tree in RAxMl with 500 bootstraps to depict the distribution/representation of the different topotypes in our data and to visualize relatedness of the serum/vaccine candidates cross-reacted in VNT assays to generate r_1_ values. Additionally, we performed a Spearman’s correlation analysis between p-amino acid distance and r1 scores. The r_1_ values were then dichotomized at 0.3, which is the threshold recommended by WHOA (22).

### Training and testing machine learning models

Using a stepwise approach, we developed a random forest classification model with sub-sampling steps and tenfold cross validation to achieve the best model performance. Upon dichotomization of the r_1_ values into binomial data, there was an imbalance in the 0 (non-cross-reacting) vs 1 (cross-reacting) classes with more than twice as many observations in the 0 category as those in the 1 category (0 = 87 observations, 1 = 21 observations). As such, there was need to subsample the data to adjust for this imbalance of the outcome in the training dataset; we thus used the synthetic minority oversampling technique (SMOTE) which statistically increased the number of cases in the dataset to balance the distribution of cases vs non-cases (1 vs 0) in the outcome to achieve better model performance. Feature-reduction was then implemented to reduce the number of model features (i.e., predictors) using the Boruta algorithm to optimize model fitting. Upon concatenating all parts of the data into a single data frame, the final data frame had 216 columns (amino acid distance and 214 site-wise amino acid differences and potential sites for N-glycosylation). With highly dimensional data, machine learning models may struggle to achieve good model accuracy and prediction due to noise introduced as the model tries to optimize the estimated contribution of each model feature to the variation in the outcome (r_1_ class in our study). As such, the Boruta algorithm has been proposed as a way of eliminating correlated and redundant features from the model, thus reducing the number of model features needed in the final model and optimizing model performance. After parsing the data through Boruta, we included all model features classified as important or tentatively important, resulting in a data frame with 35 model features. These features were VP1 amino acid distance and site-wise differences in amino acid positions (4, 13, 24, 32, 33, 43, 48, 49, 57, 69, 97, 100, 124, 135, 139-141, 143, 145, 149-151, 154, 156, 159, 166, 173, 175, 195, 198, 199, 210, 213, 214) and potential N-glycosylation profile.

Subsequently we used the 80:20 split rule to randomly split the data into three subsets to generate one training and two test datasets with 70, 17 and 21 observations each. Using *caret* (35) and *randomForest* (36) packages in R for model building, training and evaluation, including hyper-parameter tuning and 10-fold cross-validation, we trained the model on500 trees, tested it, and made predictions as summarized in the results section. We based model performance on accuracy (overall percent of observations correctly classified as cross-reacting/non-cross-reacting), sensitivity (percent of high observations correctly classified), specificity (percent of low observations correctly classified), and predictive values (proportion (%) of times the classification (non-cross-protecting-0 vs cross-protecting-1) is the true r_1_ group class.

Additionally, from the random forest model, model features were ranked in their importance to the performance of our model using mean decrease in model accuracy when a features’ data were randomized relative to the outcome (the relative prediction strength of a variable) and improvement in Gini index (measure of node impurity associated with a variable) when data were split on a variable (37, 38). This allowed us to highlight highly ranked amino acids in our VP1data, and considered the relative importance of different amino acid sites based on their role in improving model accuracy and node purity in outcome classification (15, 39).

To test the performance of the machine learning algorithm’s predictions on a completely external set of sequences that have been reported to be antigenically novel, we used data from Bachanek-Bankowska et al., (40). In this study, they reported the isolation of three antigenically distinct viruses isolated from outbreaks in Pakistan in 2016 and 2017. Bachanek-Bankowska et al., described three isolates (GenBank accession numbers MH784403, MH784404, and MH784405) from a single genetic sublineage displaying distinct antigenic phenotypes against three commonly used vaccines in the region (O 3039 and O Manisa [Boehringer Ingelheim] and O TUR 5/2009 [MSD]).

## Results

### Descriptive analysis

In this dataset, there were108 observations (serum-virus pairs) from 73 distinct viruses for which r_1_ values had been reported. In summary, the mean r_1_ value was 0.22 ± 0.23 (SD), with a range of (0.0-0.94) and a median of 0.16. Upon dichotomizing the data using the 0.3 threshold as cutoff, the two resulting groups used in subsequent modelling comprised of 87 pairs in the non-cross-neutralizing group (< 0.3 r_1_ values) and 21 in the cross-neutralizing group (≥ 0.3 r_1_ values).

Additionally, upon phylogenetic assessment, our dataset included viruses belonging to five topotypes for FMDV serotype O and the serum/vaccines used in the assays were evenly distributed among the topotype groupings (Figure 1). The mean VP1 p-amino acid distance for the pairs was 0.12 ± 0.03 with a range of 0.04 – 0.15 and median of 0.13. There was a significant (p=0.0001) moderate negative correlation (Spearman rho = -0.4) between amino acid distance and r_1_ values between the pairs (Figure 2).

**Figure 1:**
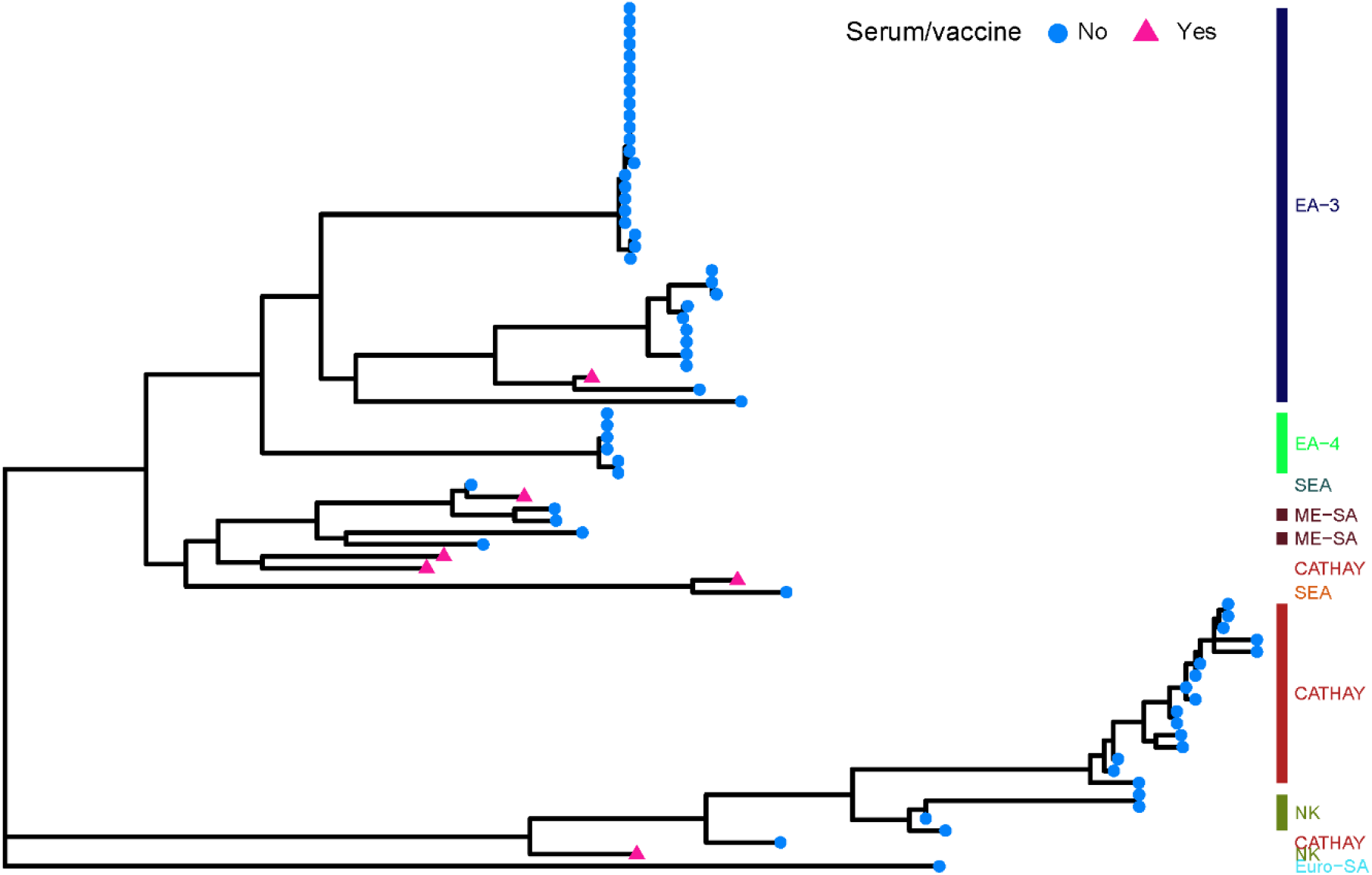
Maximum likelihood tree depicting the phylogenetic distribution of 73 foot and mouth viruses (serum, vaccine (pink triangles) and field strains (blue circles)) representing serotype O topotypes used in the development of an r_1_ predictive model. The colored bars on the right of the tree represent the different topotypes included in the data (*NK=Not Known*).

**Figure 2:**
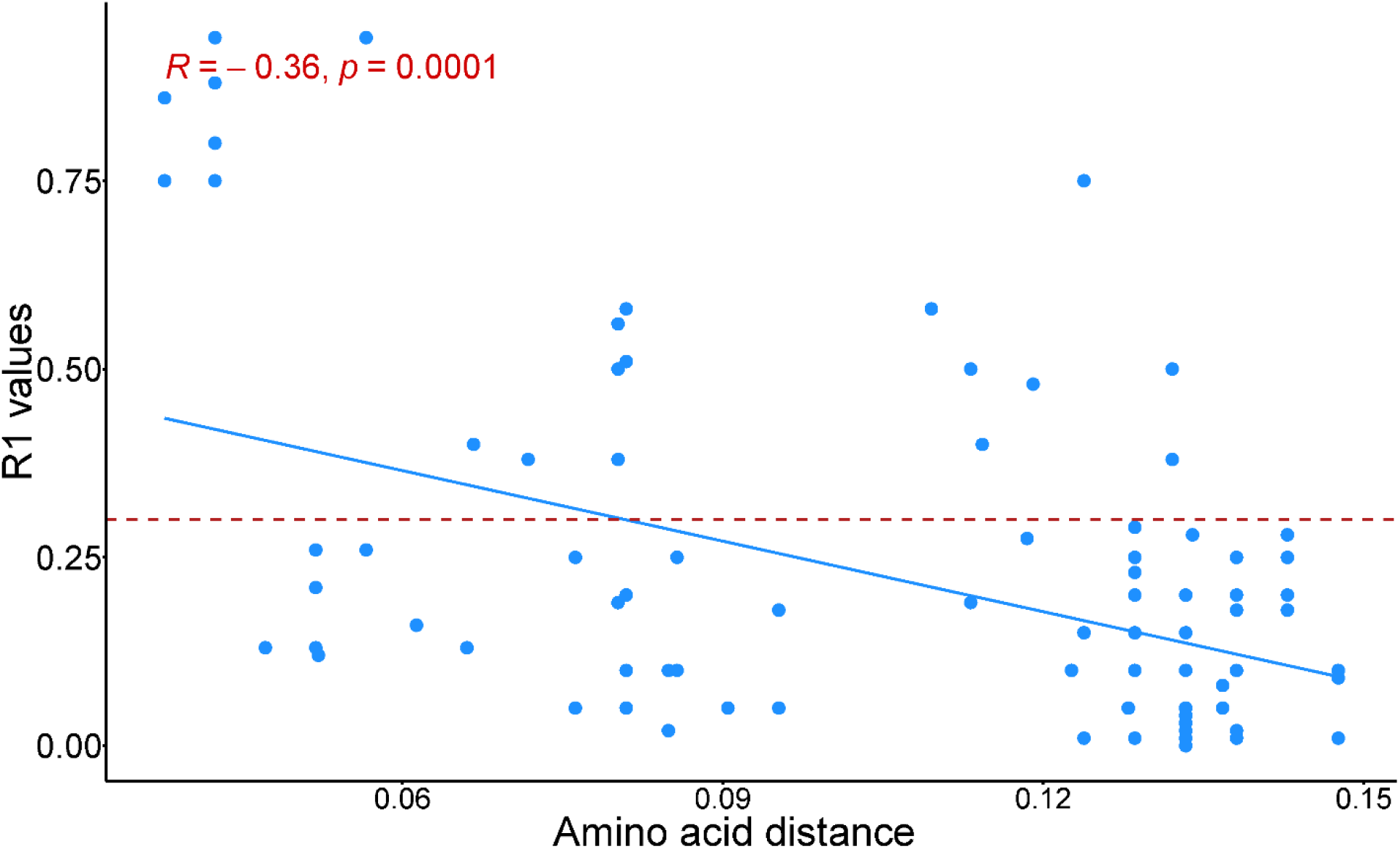
Correlation plot depicting an inverse moderate Spearman’s correlation (blue) between amino acid distance of the VP1 region for serotype O foot and mouth disease viruses and r_1_ values obtained from *in vitro* virus neutralization assays. The red-dotted line represents the cut-off of 0.3 threshold for potentials cross-neutralization between serum-virus pairs.

### Random forest model performance and predictions

The best performing model had an accuracy of 0.96 (95% CI 0.88-0.99), AUC of 0.95, sensitivity and specificity of 0.93 and 0.96, and negative and positive predictive values of 0.98 and 0.87, respectively on training data (n=70: 0=56, 1=14). On one testing dataset (n=17: 0=14,1=3), the model accuracy was 0.88 (95% CI 0.64-0.99), AUC of 0.93, sensitivity of 1.00 and specificity of 0.86, and positive and negative predictive values of 0.60 and 1.00, respectively. The model performance on the second test/validation data (n=21: 0=17, 1=4) was 1.00 accuracy (95% CI 0.84-1.00), AUC of 1.00, sensitivity and specificity of 1.00 and 1.00, and positive and negative predictive values of 1.00 and 1.00, respectively.

From the variable importance analysis that extracted which predictors had the most influence in the models’ predictive ability and accuracy, several sites were ranked highly besides the amino acid distance. These include amino acid positions 48, 100, 135, 150, and 151 in the VP1 region (Figure 3).

**Figure 3:**
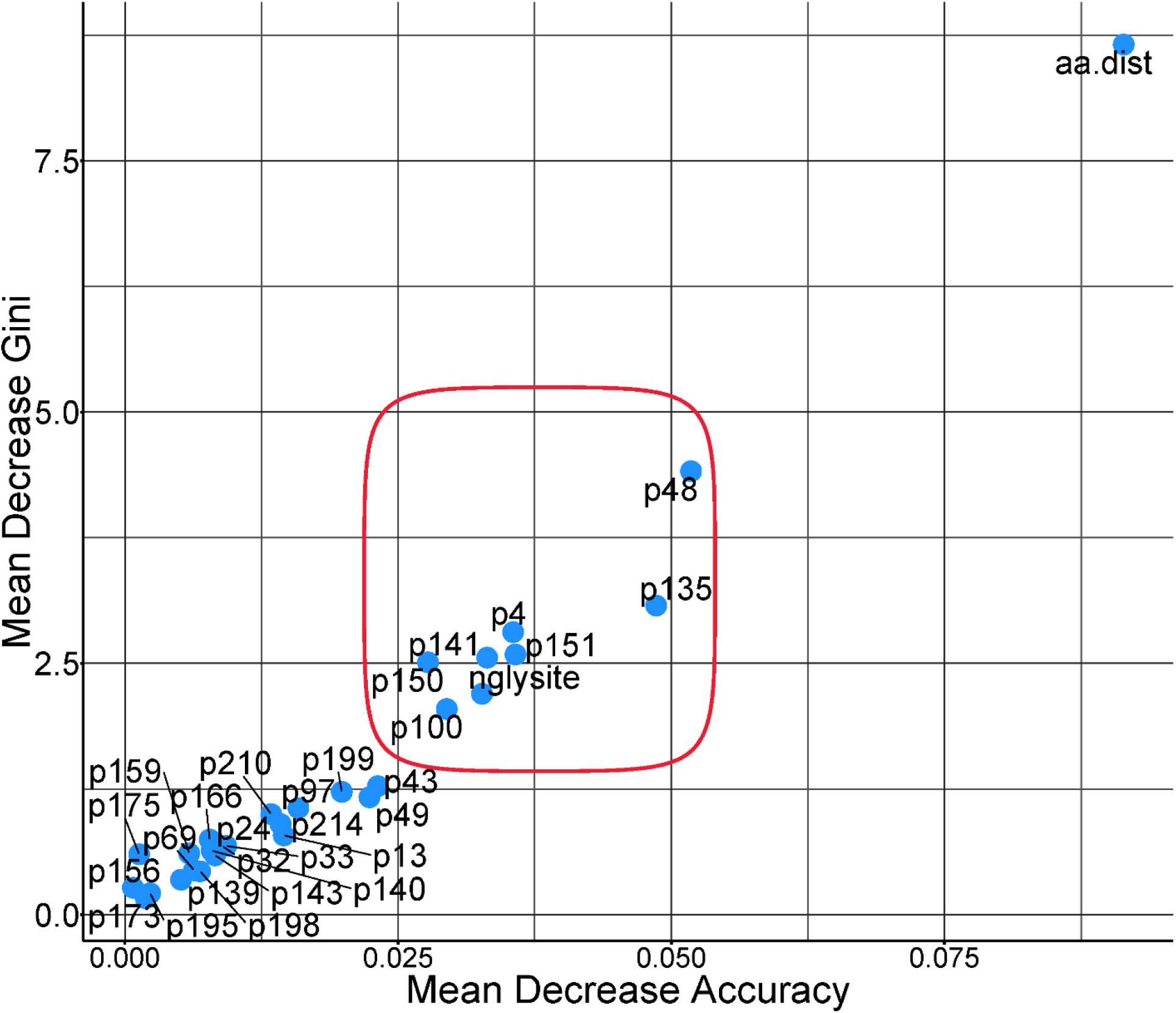
Multiway importance plot for model covariates included in the final random forest model predicting r_1_ values in VP1 region for serotype O foot and mouth disease virus. Feature labels in the graph indicate the specific amino acid site (p) while ‘aa.dist’ indicates the amino acid distance between serum-virus pairs, ‘nglysite’ similarity or difference in positions for potential N-glycosylation.

When the trained model was applied to the antigenically novel sequences described by Bachanek-Bankowska (40), the model correctly predicted the antigenic relationship of one out of three antigenically distinct viruses, i.e., between MH784405.1 (PAK/14/2017) and commonly used vaccines in the region. The model however incorrectly predicted that commonly used vaccines in the regions were good matches for both MH784403.1 (PAK/10/2016) and MH784404.1 (PAK/4/2017) (r_1_≥0.3). The *in vitro* vaccine matching experiment had indicated non-cross-neutralization between the two outbreak viruses and commonly used vaccines in this region (Supplementary table 1).

## Discussion

We developed a parsimonious random forest classification model that estimates potential cross-neutralization between serum and viruses i.e., r_1_ values for viruses belonging to serotype O FMDV with an accuracy of more than 85%. This model adds to a growing body of research and efforts towards leveraging bioinformatic data to streamline and enhance our understanding on viral antigenicity and host immune response as exemplified in diseases like influenza (16, 17, 41–44). diseases like influenza. Optimized and robust predictive models can be valuable in decision making and implementing strategic interventions towards the control, and ultimately eradication, of FMDV. This model requires further training and refinement with diverse datasets because when applied to the antigenically novel sequences described by Bachanek-Bankowska (40), the model correctly predicted the antigenic relationship of only one of three novel viruses.

The suitability of using *in vitro* methods to estimate *in vivo* cross-protection in FMD (45), r_1_ values and 0.3 thresholds compared to other measures of immune response and cross reactivity has been argued by several studies (14, 45–48). However, though this model is based on r_1_ values it can be modified to accommodate other measures of antigenic variability like raw VNT results. Since r_1_ values are still the most-used option when conducting vaccine matching and selection of candidates, whether achieved through simple ELISA or a modified combination of techniques to estimate cross-protection, our model is well trained to aid in the identification of potential cross-neutralizing and non-cross-neutralizing sera thus streamlining the selection of potential vaccine candidates and facilitating immediate comparisons and of field strains where necessary. Moreover, when applied in non-research settings, animal health agencies/institutions can easily upload sequences to the model from isolates or clinical samples and promptly identify potential options for immunization. The continued improvement and accessibility of sequencing technology such as Minion Nanopore sequencing, which can be deployed in LMIC settings, may also increase the frequency and capacity of in-country generation of sequences (especially in countries where access to reference labs is limited) complementing the utility of predictive tools supported by this model.

By highlighting the highly ranked/important amino acids in the VP1 region, we posit that those amino acids influence host immune induction and response as differences in those sites appear to be important for estimating the r_1_ values between serum-virus pairs. This supports findings by other studies (10, 12) that certain immunodominant sites exist in the VP1 region. To better understand challenges with vaccine effectiveness, causes of vaccine failure and breakthrough outbreaks in vaccinated populations as well as the mechanisms of emergence and spread of new viruses, dynamics of mutations in these sites can be investigated further (49). In these data, seven amino acid positions were most influential in the ability of our model to accurately distinguish between cross-neutralizing (≥0.3) vs non-cross-neutralizing (<0.3). These were amino acids in positions 4, 48, 100, 135, 141, 150 and 151. Except position 4 and 100, all these amino acids are part of immunogenically important regions described as the B-C and G-H loops in the VP1 protein of serotype O (6) in which certain mutations in combination or independently have been thought to confer serological heterogeneity among FMDVs (50). In sequences used for this study, N-glycosylation was predicted to occur most often at sites 86, then 101, and least often at site 132.

Even with improved predictive ability to accurately classify cross-neutralizing sera/vaccine as possible, there are drawbacks of *in silico* modelling for biological processes. The immune response is a composite process which commonly involves the interplay of multiple compartments of the host immune response system and different factors (e.g., environment and host genetics) that may influence host-virus interactions. Such factors contribute to why measures of cross-reactivity *in vitro* often do not translate well to *in vivo* cross-protection in the field. Our model does not account for these factors. Nevertheless, with current data, we were able to accurately categorize 8 to 9 out of every 10 serum-virus pairs for serotype O as cross-neutralizing or non-cross-neutralizing (based on r_1_ values). However, the wider confidence interval suggests that there is some misclassification and misidentification of sera/vaccine candidates.

As such, continued improvement of *in silico* models is necessary both with diverse datasets and potential validation with *in vivo*/field application data as attempted in this study. That limitation notwithstanding, with the limited data used here, the model can at least aid in filtering potential candidates to be considered for in-depth *in vitro*/*in vivo* assays or blends in vaccination programs which would save resources and increase efficiency of the decision-making process and FMD management.

Lastly, other VP regions in the P1 portion of the genome also likely play a role in host-virus interaction and immune response. However, since we only relied on publicly available secondary data, we did not have access to full genome sequences for all 73 viruses included in the study hence the need to restrict the analysis to the role of VP1 in estimating r_1_ values. Whole genomic data could improve our model. Future efforts will explore the benefits of a broader genomic analysis and more sensitive threshold of cross-protection.

## Conclusion

Ultimately, this study adds to the growing body of literature that machine learning can be applied to genetic/genomic data to achieve a more nuanced analysis of the relationship between genetic variability and cross-recognition of viruses by host immune systems. Such models can support immunization as a pathway to disease management. In this study, we demonstrate the opportunities and potential for leveraging routinely generated sequence data and the application of a parsimonious machine learning model to streamline the process of decision-making in vaccine development and application to control FMD, especially serotype O (but also applicable to other serotypes). Specifically, by obtaining an accurate model with high sensitivity and specificity, appropriate vaccine candidates may be able to be selected more quickly, although *in vivo* experiments would still ultimately be necessary to assess cross-protection. Also, this capability would enable tailoring immunization protocols for use in the field with a faster turnaround time for decision-making. Lastly, outputs from such a model can be combined with other mathematical models to understand drivers and trends of viral emergence, especially the role of immune pressure in driving the evolution and spread of FMDV. The latest version of the r_1_ predictive model is available for access via a Shiny dashboard (https://dmakau.shinyapps.io/PredImmune-FMD/).

## Data availability

The nucleotide sequences of the FMDV used in this analysis are already publicly available on GenBank and the list provided in Supplemental material 1 can help in identification and downloading of the sequences if needed.

## Supplemental material

All supplementary materials have been provided in one file.

## Acknowledgements

This project was supported by the USDA Agricultural Research Service, award 58-8064-2-006. The authors declare there are no conflicts of interest.

