## Supplemental material 1 for "Machine learning approaches for estimating cross-neutralization potential among FMD serotype O viruses"

**List of accession numbers of virus and vaccine/serum isolates included in this study. All viruses were obtained from studies by these 4 studies** (Maree et al., 2011; Tesfaye et al., 2020; Upadhyaya, Mahapatra, Mioulet, & Parida, 2021; Yang, Xu, Goolia, & Zhang, 2014)

KJ831676.1,DQ164880.1,DQ164881.1,KM243162.1,KM243163.1,MZ851288.1,MZ851289.1,MZ851290.1,MZ851291.1,MZ851292.1,MZ851293.1,MZ851294.1,MZ851295.1,MZ851296.1,MZ851297.1,MZ851298.1,MZ851299.1,MZ851300.1,MZ851301.1,MZ851302.1,MZ851303.1,KJ831676.1,DQ164880.1,DQ164881.1,KM243162.1,KM243163.1,MZ851288.1,MZ851289.1,MZ851290.1,MZ851291.1,MZ851292.1,MZ851293.1,MZ851294.1,MZ851295.1,MZ851296.1,MZ851297.1,MZ851298.1,MZ851299.1,MZ851300.1,MZ851301.1,MZ851302.1,MZ851303.1,DQ164880.1,DQ164881.1,KM243162.1,KM243163.1,MZ851288.1,MZ851289.1,MZ851290.1,MZ851291.1,MZ851292.1,MZ851293.1,MZ851294.1,MZ851295.1,MZ851296.1,MZ851297.1,MZ851298.1,MZ851299.1,MZ851300.1,MZ851301.1,MZ851302.1,MZ851303.1,KJ831676.1,DQ164880.1,DQ164881.1,KM243162.1,KM243163.1,MZ851288.1,MZ851289.1,MZ851290.1,MZ851291.1,MZ851292.1,MZ851293.1,MZ851294.1,MZ851295.1,MZ851296.1,MZ851297.1,MZ851298.1,MZ851299.1,MZ851300.1,MZ851301.1,MZ851302.1,MZ851303.1,KJ606977.1,KJ606980.1,KJ606981.1,KJ606983.1,KJ606984.1,KJ606978.1,KC519630.1,KJ606979.1,KJ606982.1,MN518164.1,MN518166.1,MN518152.1,MN518153.1,MN518154.1,MN518157.1,MN518149.1,MN518151.1,MN518142.1,MN518146.1,MN518147.1,MN518143.1,MN518155.1,MN518144.1,MN518145.1,MN518148.1.

| Outbreak strain | Commonly used vaccines | Predicted $r_1$ class (1= ≥0.3) |
|---|---|---|
| AY593823.1o1manisaiso87 | JF968170.1HongKong/3039/2004 | 1 |
| AY593823.1o1manisaiso87 | MT443823.1TUR/5/2009 | 1 |
| AY593823.1o1manisaiso87 | MH784405.1_PAK/14/2017 | 1 |
| AY593823.1o1manisaiso87 | MH784403.1_PAK/10/2016 | 1 |
| AY593823.1o1manisaiso87 | MH784404.1_PAK/4/2017 | 1 |
| JF968170.1HongKong/3039/2004 | AY593823.1o1manisaiso87 | 1 |
| JF968170.1HongKong/3039/2004 | MT443823.1TUR/5/2009 | 0 |
| JF968170.1HongKong/3039/2004 | MH784405.1_PAK/14/2017 | 1 |
| JF968170.1HongKong/3039/2004 | MH784403.1_PAK/10/2016 | 1 |
| JF968170.1HongKong/3039/2004 | MH784404.1_PAK/4/2017 | 1 |
| MH784403.1_PAK/10/2016 | JF968170.1HongKong/3039/2004 | 1 |
| MH784403.1_PAK/10/2016 | AY593823.1o1manisaiso87 | 1 |
| MH784403.1_PAK/10/2016 | MT443823.1TUR/5/2009 | 1 |
| MH784403.1_PAK/10/2016 | MH784405.1_PAK/14/2017 | 1 |
| MH784403.1_PAK/10/2016 | MH784404.1_PAK/4/2017 | 1 |
| MH784404.1_PAK/4/2017 | JF968170.1HongKong/3039/2004 | 1 |
| MH784404.1_PAK/4/2017 | AY593823.1o1manisaiso87 | 1 |
| MH784404.1_PAK/4/2017 | MT443823.1TUR/5/2009 | 1 |
| MH784404.1_PAK/4/2017 | MH784405.1_PAK/14/2017 | 1 |
| MH784404.1_PAK/4/2017 | MH784403.1_PAK/10/2016 | 1 |
| MH784405.1_PAK/14/2017 | JF968170.1HongKong/3039/2004 | 1 |

| | | |
|---|---|---|
| MH784405.1_PAK/14/2017 | AY593823.1o1manisaiso87 | 1 |
| MH784405.1_PAK/14/2017 | MT443823.1TUR/5/2009 | 1 |
| MH784405.1_PAK/14/2017 | MH784403.1_PAK/10/2016 | 1 |
| MH784405.1_PAK/14/2017 | MH784404.1_PAK/4/2017 | 1 |
| MT443823.1TUR/5/2009 | JF968170.1HongKong/3039/2004 | 0 |
| MT443823.1TUR/5/2009 | AY593823.1o1manisaiso87 | 1 |
| MT443823.1TUR/5/2009 | MH784405.1_PAK/14/2017 | 1 |
| MT443823.1TUR/5/2009 | MH784403.1_PAK/10/2016 | 1 |
| MT443823.1TUR/5/2009 | MH784404.1_PAK/4/2017 | 1 |

**Table 1: Predicted cross-reaction classification between 3 outbreak strains reported in Pakistan between 2016-2017 and commonly used vaccines in the region used in a vaccine matching experiment as described by Bachanek-Bankowska** (Bachanek-Bankowska et al., 2019)
